# Ancient Roman saltworks drive the present-day microbial community profiles in a coastal aquifer

**DOI:** 10.64898/2025.12.23.695991

**Authors:** M. Boccanera, S. Amalfitano, M. Melita, D. Corso, A. Valle, S. Ghergo, E. Preziosi

## Abstract

Historical salinization from ancient saltworks can leave a long-lasting imprint on coastal aquifers, but its impacts on subsurface microbial communities and ecosystem functioning remain poorly understood. This study examined how legacy salt inputs from Roman saltworks in the Tiber River delta (Fiumicino, Italy) continue to shape present-day groundwater chemistry, microbial community structure, and metabolic potential of the coastal aquifer. We analyzed groundwater samples from non-salinized and salinized units of the same aquifer using integrated hydrogeochemical characterization, flow cytometry, 16S rRNA gene amplicon sequencing, and functional metabolic assays. Salinized samples exhibited elevated chloride, bromide, sodium, and sulfate concentrations, with distinctive ionic ratios (Br/Sr, Cl/K, SO₄/Ca) indicating dissolution of salt deposits rather than contemporary seawater intrusion. Salinization reduced the microbial diversity and shifted communities from diverse freshwater-adapted families toward an abundant halotolerant assemblage dominated by Campylobacterota (families Sulfurimonadaceae and Sulfurovaceae). Functional annotation suggested broadly conserved potentials for carbon, nitrogen, and sulfur cycling.

However, the Biolog assays revealed higher heterotrophic respiration and carbon substrate use but lower functional diversity in salinized samples, with particularly enhanced polymer degradation. Ordination analyses showed a clear separation of aquifer units along the salinization gradient, with coordinated chemical and microbial vectors indicating alternative ecosystem states sustained by millennia-old anthropogenic salt inputs. Our findings showed that ancient saltworks can drive persistent hydrogeochemical alteration, select specialized halotolerant microbiomes, and reconfigure carbon and nutrient processing while maintaining core biogeochemical functions, with critical implications for coastal groundwater management strategies.

## 1. Introduction

Groundwater salinization is recognized as a major threat to water resources since the accumulation of salts in underground environments, particularly sodium and chloride ions, can directly affect the aquifer chemical composition and the ecosystem functioning, limiting the suitability of groundwater resources for drinking, irrigation, and industrial uses (Foster et al., 2018; Mueller et al., 2024; Ghirardelli et al., 2025). Land use and local hydrogeological conditions also influence the spatial distribution and level of salinity within the aquifer, with direct effects on all groundwater-dependent ecosystems, such as wetlands, marshes, and rivers (Bierkens and Wada, 2019).

In coastal settings, salinization often results from seawater intrusion, the movement of seawater into zones previously occupied by fresh groundwater. Excessive groundwater extraction frequently intensifies this process, although it can occur naturally (Werner et al., 2013; Jasechko et al., 2020). While much attention has focused on seawater intrusion, it is now recognized that legacy salt sources, such as paleointrusions, relic saline deposits from ancient evaporative environments, and historic saltwork activities, can contribute to groundwater salinity, independently of current seawater contact, suggesting complex histories of sedimentary salt accumulation and transformation (Li et al., 2020; Dang et al., 2022; Cui et al., 2025). These salinity hotspots, located far inland from the coastlines, are visible in the unique biodiversity and ongoing geochemical processes but can pose contemporary issues for groundwater resource management (Khalil et al., 2025; Passarella et al., 2025). Historic saltworks have held critical significance, shaping coastal landscapes and ecosystems worldwide.

While Mediterranean saltworks are often cited for their scale and antiquity (Mastrorillo et al., 2016; Marien et al., 2023), large-scale salt production developed in China, India, Central America, and Africa, with ancient societies building evaporation basins, salinas, and processing facilities along coasts (Flad, 2007; Mani et al., 2012; Antonites, 2013; Watson and McKillop, 2019). In addition to significant cultural and economic effects, supporting local economies, international trade, food preservation across civilizations, major habitat alterations, and aquifer salinization are part of their lasting legacy (Harding, 2014; Marzano, 2024).

Within saline aquifers, microbial communities perform essential ecosystem functions, driving biogeochemical cycling and nutrient transformation (Ma et al., 2022; Sang et al., 2025). Therefore, alterations in microbial community profiles will promptly reflect on groundwater chemistry, quality, and ecosystem services (Houghton et al., 2023). In particular, salinization can impose a strong selective pressure, since only halotolerant and halophilic microbes can persist, generally resulting in communities with reduced species richness and evenness (Chen et al., 2024; Nelson et al., 2024). At the phylum level, Proteobacteria and Bacteroidetes typically dominate across all salinity levels, but finer taxonomic resolution revealed that members of the class Gammaproteobacteria (including the order Alteromonadales, the family Alteromonadaceae and the genus *Marinobacter*) were increasingly abundant in more saline waters (Liu et al., 2018; Gorrasi et al., 2021; Zhi et al., 2024). Functional gene surveys further indicated an enrichment of genes associated with dissimilatory sulfate reduction under higher salinity, suggesting enhanced potential for sulfur and carbon cycling in salt-impacted zones (Li et al., 2024; Sang et al., 2025). Other studies reported that overall microbial diversity may remain resilient, and key functional taxa are maintained despite high salinity, emphasizing the importance of the geological sediment type and the water origin (Rath and Rousk, 2015; Zhang et al., 2021).

Despite significant research advances, major knowledge gaps remain in understanding how historical salinization processes shape aquifer microbial communities, restructure ecological functions, and affect the present-day ecosystem services (Seibert et al., 2024). Moreover, it remains unclear whether halotolerant microorganisms can fully compensate for the ecological roles of microbial taxa that decline under elevated salinity (Nelson et al., 2024). Broader investigation across diverse geological settings and supporting field data on microbial community shifts across salinity gradients is required to predict the resilience of groundwater ecosystems and to inform sustainable management of coastal groundwater resources (Wang et al., 2025).

The objectives of this study were to (i) explore whether the microbial community profiles changed in non-salinized and salinized units of the same aquifer, (ii) identify specific halotolerant taxa that may become dominant following historical salinization, and (iii) investigate how salinization-induced microbial community shifts can influence groundwater biochemical processes and nutrient cycling. We hypothesized that historical salinization of coastal aquifers can reshape the present-day microbial communities by altering their structure and function in ways that consequently impact aquifer ecology and ecosystem services.

## 2. Materials and methods

### 2.1 Study area and hydrogeochemical setting

The study was conducted within the municipality of Fiumicino (Lazio, Italy) at the north bank of the Tiber River, an area encompassing a coastal aquifer of considerable importance for irrigation and other local applications (Figure 1). During the upper Pleistocene and Holocene, and until the 19^th^ century, the area was characterized by extensive marshlands responsible for the accumulation of peat-rich sediments with medium to low permeability. In historical times, these marshes hosted large saltworks exploited by the Etruscans and Romans for salt extraction (Morelli and Forte, 2014). The Late Quaternary Tiber River depositional sequence (“Tiber River Synthem” of the official Geological Map of Italy at 1:50,000 scale) (Funiciello and Giordano, 2008) was extensively investigated and reconstructed based on the stratigraphy of available cores (Milli et al., 2013). A complex interplay of tectonic uplift, volcanic activity, and glacio-eustatic sea-level fluctuations accounts for the vertical and lateral heterogeneity of the stratigraphy. Specifically, around 11,550 yr BP during the transition from the Upper Pleistocene to the Holocene, a line of barrier islands developed along the coastline during a transgressive phase from lowstand to highstand conditions.

**Figure 1.**
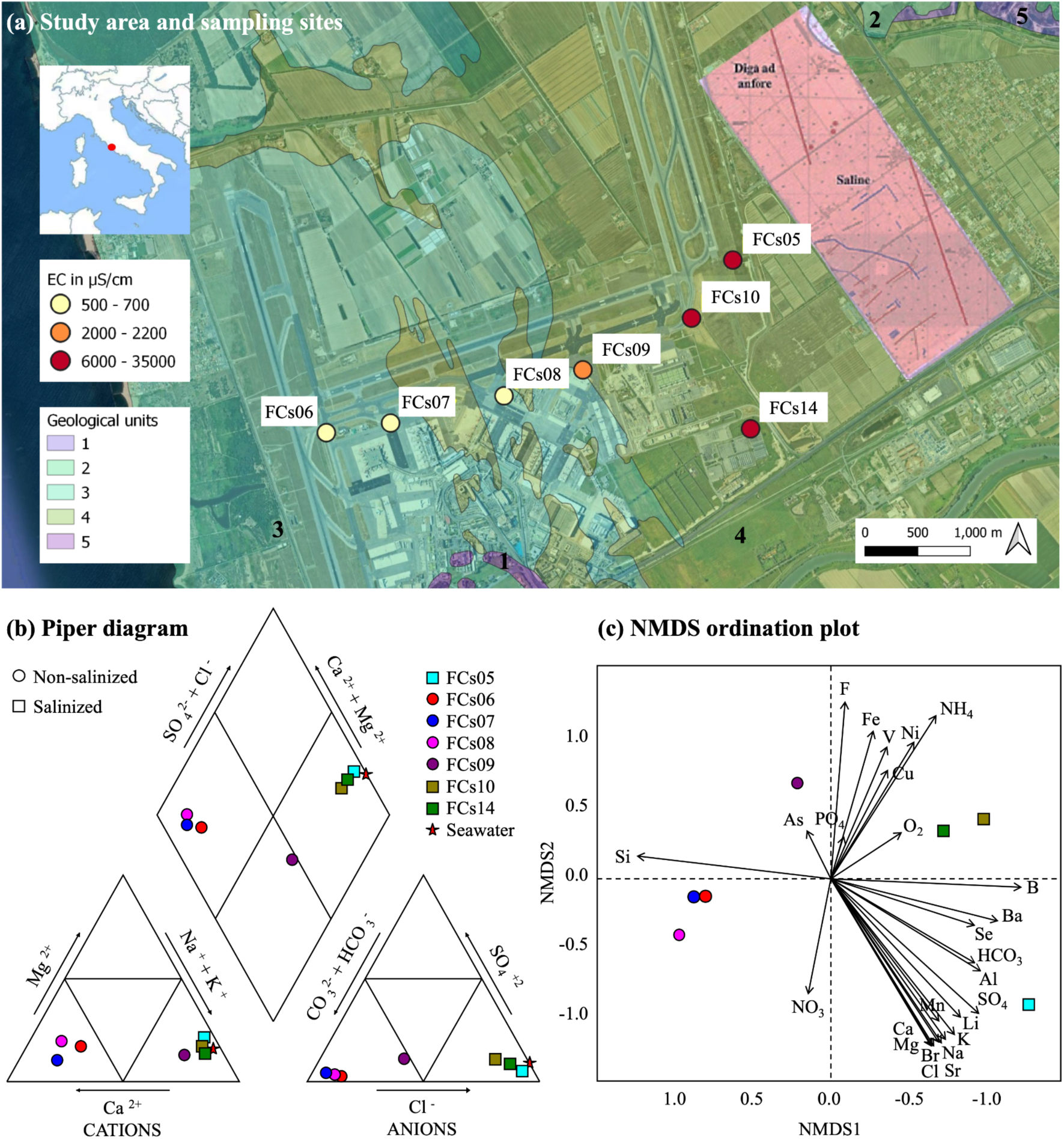
(a) Aerial view of the study area showing the sampling sites, the ancient Roman saltworks (pink area as presented in Morelli et al., 2014), and the major geological units (1 = anthropogenic deposits (Holocene); 2 = eolian sands and dunes (Upper Pleistocene-Holocene); 3 = fluvio-lacustrine and marsh sediments rich in peat (Upper Pleistocene-Holocene); 4 = Ponte Galeria formation (Middle Pleistocene). (b) The Piper diagram illustrates the primary ionic composition of the groundwater samples. The standard seawater was included as a saline endmember. (c) The NMDS ordination plot represents the chemical composition of groundwater samples, including all assessed major compounds and trace elements.

Brackish lagoons formed landward of the barrier system, together with marshes where organic-rich fine sediments and peat accumulated. Starting around 6,000 yr BP, beach ridges prograded seaward (westward) in front of the marshes and lagoons. A sedimentary succession 70–80 m thick (comprising fluvial and fluvio-lacustrine deposits, marsh sediments rich in peat, beach sands of Upper Pleistocene-Holocene, and coastal-transitional alluvial sediments of the Middle Pleistocene Ponte Galeria formation) overlies the Lower Pleistocene marine clays of the Monte Mario Formation across an erosional surface formed during a sea-level fall (Valle, 2023). As a result of this evolutionary history, the eastern sector of the area is dominated by organic-rich fine sediments, whereas eolian sands prevail toward the west. The water chemistry of the target area relies on three interactive processes, including (i) the water-rock interaction with eolian sands (West) and palustrine-lacustrine deposits (East), (ii) the contribution of marine water via post-glacial sea-level variations and the Holocene brackish ponds, and (iii) the natural and induced evaporation of saline water (saltworks) in historical time (Morelli and Forte, 2014; Mastrorillo et al., 2016).

### 2.2. Field sampling and physical-chemical analyses

Groundwater was sampled in June 2023 from seven piezometers. Purging (> 30 min) was performed with a low-flow submersible pump (Proactive Mega-Monsoon 12V Plastic Pump connected to a Low Flow Power Booster 3 Controller) until stabilization of physical-chemical parameters, using a multiparametric flow-cell probe (Royal Eijkelkamp Scuba 50 multiprobe) to measure temperature (T, °C), pH, oxidation-reduction potential (ORP, mV), electrical conductivity (EC, mV), and dissolved oxygen (DO, mg/l). Groundwater samples were filtered through 0.45-µm pore-sized HNO₃ 1% treated polycarbonate filters, collected in HNO₃ 1% treated polyethylene bottles, and stored at 4°C until analysis. One fraction was acidified to pH < 2 (1% HNO₃) immediately after filtration for assessing the concentrations of major cations and trace elements.

Samples were analyzed for anions within 24 hours from sampling by means of Ionic Chromatography (IC, Dionex DX-120), while bicarbonates were quantified through HCl titration of 50 mL of filtered sample using a HACH-Lange Titralab AT1000. The acidified aliquot was analyzed in Optical Emission Spectroscopy (ICP-OES) for major cations and Inductively Coupled Plasma Mass Spectrometry (ICP-MS, Agilent Technologies 7500c) for trace elements. The overall quality of analysis is tested by the electro-neutrality (EN% < 5%), calculated using the equation (Appelo and Postma, 2004):

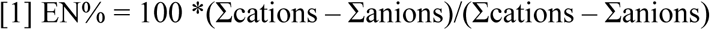

Certified materials (NIST 1640a, Trace elements in natural waters) were used to check the quality of the laboratory results for trace elements. Non-detectable values were replaced by half of their detection limits (USEPA, 2000).

### 2.3. Molar ion ratios and saline water fraction

Total dissolved solids (TDS, mg/l) were estimated analytically as the sum of the concentrations of the major dissolved ions quantified in each groundwater sample. The molar ratios between major hydrochemical ions (expressed as meq/L) were computed to evaluate salinization processes, water-rock interactions, potential solute origins, and the geochemical evolution of the coastal aquifer (Han and Currell, 2018). To maintain consistent scaling and facilitate comparison among samples, the average concentration of the ion at higher molarity was divided by that at lower molarity.

The fraction of saline water in the samples was calculated following the equation:

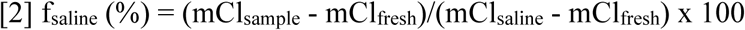

where m is the molar concentration (Jiao and Post, 2019).

The lowest and highest Cl concentrations registered in the target aquifer served as endmember values for local fresh and saline groundwater, respectively. Samples with a low saline water mixing fraction (f_saline_ < 1%) were considered representative of the non-salinized aquifer unit (i.e., FCs06, FCs07, FCs08, and FCs09), while samples with higher values (f_saline_ > 1%) were identified as representative of the salinized aquifer unit (i.e., FCs05, FCs10, and FCs14).

### 2.4. Microbial load assessment by flow cytometry

The microbial load of each groundwater sample was assessed by the flow cytometer A50-micro (Apogee Flow System, Hertfordshire, UK), equipped with a 20 mW solid-state blue laser (488 nm) and a 16 mW solid-state UV laser (375 nm). Water aliquots were immediately placed in 2-mL safe-lock tubes, fixed with formaldehyde (2% final concentration), stored at 4°C, and analyzed the day after sampling. The light scattering signals (forward and side light scatter named FSC and SSC, respectively), red fluorescence (>610 nm), orange fluorescence (590/35 nm), and blue fluorescence (430–470 nm) were acquired and considered for the direct identification and quantification of distinct microbial groups by following harmonized protocols (Gasol and Morán, 2015). Data handling and visualization were performed by the Apogee Histogram Software (v89.0). The total cell counts (TCC) were determined by the signatures in a plot of the side scatter vs. the green fluorescence. The intensity of green fluorescence emitted by SYBR-positive cells allowed the discrimination between two cell groups with low (LNA) and high (HNA) nucleic acid content (Amalfitano et al., 2018).

### 2.5. Phylogenetic community composition and functional annotation by 16S rRNA gene amplicon sequencing

Water samples (0.5 - 3 L) were vacuum filtered on the same day using polycarbonate filter membranes (type GTTP; pore size = 0.2 µm; diameter = 47 mm; Millipore, Eschborn, Germany). DNA extraction was performed from filter membranes using the DNeasy PowerSoil Pro kit (QIAGEN, Hilden, Germany) and following the manufacturer’s protocol. The NanoDrop ND-1000 spectrophotometer (NanoDrop Technologies, Wilmington, DE, USA) was used to check the DNA’s concentration and purity. The extracts were then stored at −80°C until they were ready for further analysis. PCR amplification of the V3-V4 hypervariable region of the 16S rRNA gene was performed using the 341F-805R primer pair(Klindworth et al., 2013),followed by PCR product purification. Nextera library indexing and preparation for sequencing were conducted using a 2 × 300 bp paired-end protocol as described elsewhere (Corso et al., 2025). Sequencing was carried out on the Element Biosciences AVITI platform (Garbe et al., 2024). Obtained reads were processed using R (version 4.5.1) with the DADA2 pipeline designed for BigData paired-end reads (https://benjjneb.github.io/dada2/bigdata_paired.html) (Callahan et al., 2016), which enabled the identification of amplicon sequence variants (ASVs). Additionally, following the guidelines provided in the tutorial at https://github.com/nuriamw/micro4all, we applied Figaro (Weinstein et al., 2019) to set optimal trimming parameters and Cutadapt (Martin, 2011) to remove adapter or primer sequences from the raw reads. The resulting ASVs were then taxonomically annotated using the SILVA reference database (version 138.2) (Quast et al., 2012). The alpha diversity (H_phylo_) and the differential abundance analysis was performed using the R microeco package (Liu et al., 2025) to identify microbial taxa that most contributed to the differentiation among sample groups. The Linear Discriminant Analysis Effect Size (LEfSe) analysis (Segata et al., 2011) was adopted to identify microbial taxa significantly enriched in non-salinized and salinized samples (herein defined as freshwater-adapted and halotolerant taxa, respectively).

The FAPROTAX database was employed to annotate the functional metabolic capacity of ASVs according to their taxonomy (Louca et al., 2016). Normalized counts by median sequencing depth were used as input. The FAPROTAX functional data were first converted into a long-format dataframe. To facilitate ecological interpretation, each function was then assigned to the major biogeochemical cycles (i.e., carbon, nitrogen, or sulfur) according to established lists of FAPROTAX functional groupings. A bubble plot was subsequently generated using ggplot2, categorizing the samples into panels representing salinized and non-salinized aquifer units.

### 2.6. Microbial functional profiling by Biolog™ assays

Microbial heterotrophic respiration, metabolic potential, metabolic profiling, and functional diversity (H_funct_) were evaluated using Biolog™ MT2 and EcoPlates assays (Biolog, Inc., Hayward, CA, USA). Both tests consist of 96-well microplates containing the respiration-sensitive dye tetrazolium chloride, whose reduction to formazan indicates microbial degradative activity. In MT2 plates, no external substrates are supplied, and the signal depends on the degradation of organic matter naturally present in the sample, reflecting the heterotrophic respiration rates. The EcoPlates assay provides 31 organic carbon sources in triplicate, categorized into six substrate categories, including amino acids (n = 6), amines (n = 2), carbohydrates (n = 9), carboxylic acids (n = 8), phenolic compounds (n = 2), and polymers (n = 4), enabling the assessment of potential metabolic capacity, and the substrate utilization profiles (Melita et al., 2023). On the same sampling day, 125 µl of sample were inoculated in each microplate well. The initial Optical Density (OD) was recorded at 590 nm using a spectrophotometric plate reader (PerkinElmer VICTOR™ X3 Multilabel Plate Reader). The plates were incubated at 20°C for 24 h, after which OD was measured again to quantify the reduction of colorless tetrazolium chloride to purple formazan, indicative of microbial metabolic activity. In both assays, absorbance values were corrected prior to analysis by subtracting the initial (time 0) OD to account for inoculum density. The microbial heterotrophic respiration was assessed by calculating the average values of the Biolog™ MT2 test. For the Biolog™ EcoPlates tests, control well (no substrate provided) values were further subtracted to isolate substrate-specific responses (Németh et al., 2021). The metabolic potential and metabolic profiles were obtained by calculating the average well-color development (AWCD) and the average well-color development per substrate category (AWCDs) on the EcoPlates values following the formulas:

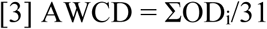

where OD_i_ is the corrected absorbance value of each well,

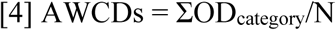

where OD_category_ is the corrected absorbance value of the substrates within the substrate category and N is the number of substrates within that category.

### 2.7. Statistical Analysis

Given the non-normal distribution of most available data, non-parametric tests were selected for assessing associations within the dataset. The Kruskal–Wallis test was used to assess differences in median values of each variable between non-salinized and salinized sample groups (p < 0.05). The multi-group SIMilarity PERcentage test (SIMPER), using the Bray-Curtis similarity measure of log-transformed data, was run to identify the major chemical components and trace elements (expressed as mg/l) that were primarily responsible for observed differences between the non-salinized and salinized aquifer units, then sorted in descending order of their contribution to the average dissimilarity (Melita et al., 2019). The one-way Analysis of Similarity (ANOSIM), using Bray-Curtis and Gower distance, was performed by applying the vegan and cluster packages to assess differences in microbial community composition between non-salinized and salinized aquifer units (Maechler et al., 1999; Oksanen et al., 2001). Spearman’s rank correlation coefficient (r_s_) was applied to assess the statistical relationships between the relative abundances of selected microbial taxa and their corresponding environmental parameters (p < 0.05). The nonmetric multidimensional scaling ordination (NMDS) analysis was performed using Bray-Curtis or Gower dissimilarity matrices. Results were visualized in a two-dimensional plot highlighting differences between non-salinized and salinized samples. All statistically different parameters, including chemical and microbial community descriptors, were incorporated in the analysis with a vector-fitting procedure, in which the length of the arrow is proportional to the correlation between the NMDS axes and each descriptor (Amalfitano et al., 2014).

## 3. Results

### 3.1. Effects of historical salinization on groundwater chemical composition

Field parameters revealed significant differences in EC and Cl levels between non-salinized samples collected from western wells (i.e., closer to the coastline) and salinized samples collected from the easternmost wells (i.e., more inland). Groundwaters were anoxic (DO = 0.2-1.5 mg/L), with redox conditions varying from -201 mV (FCs09) to +250 mV (FCs08). The pH ranged from slightly acidic to neutral, reaching a maximum of 9.7 in FCS08. (Figure 1, Supplementary Table 1).

The saline water mixing fraction was negligible (f_saline_ < 1%) in groundwater samples collected from the non-salinized aquifer unit, with a relatively higher value registered in FCs09 (f_saline_ = 0.8%). Consistently higher contributions of saline water were found in samples from the salinized aquifer units, with the highest value registered in FCs05 (f_saline_ = 38.8%). The TDS concentrations increased consistently, transitioning from fresh (< 1000 mg/l) to brackish (1000–10000 mg/l) to saline conditions (> 10000 mg/l). The salinized aquifer unit showed significantly higher levels of Cl, Br, Na, Mg, SO₄, Al, Sr, K, and Ca. Only Si was significantly more concentrated in the non-salinized aquifer unit (Table 1).

**Table 1.** Chemical components with significant differences between their median values in the non-salinized and salinized aquifer units (Kruskal–Wallis test, p < 0.05). Sampling sites are presented in ascending order of Cl levels. Chemical parameters are ordered left to right by their relative contribution to average group dissimilarity (SIMPER test).

| | Cl<br>mg/l | Br<br>mg/l | Na<br>mg/l | Mg<br>mg/l | $\text{SO}_4$<br>mg/l | Al<br>mg/l | Sr<br>mg/l | K<br>mg/l | Ca<br>mg/l | Si<br>mg/l |
| --- | --- | --- | --- | --- | --- | --- | --- | --- | --- | --- |
| **Non-salinized** |  |  |  |  |  |  |  |  |  |  |
| FCs07 | 13.6 | 0.04 | 14.3 | 8.4 | 13.0 | 0.006 | 0.28 | 12.3 | 89.7 | 11.0 |
| FCs08 | 16.6 | 0.03 | 10.6 | 11.1 | 7.4 | 0.018 | 0.26 | 4.0 | 59.4 | 9.0 |
| FCs06 | 31.1 | 0.23 | 27.8 | 13.8 | 8.1 | 0.013 | 0.22 | 7.6 | 75.8 | 10.2 |
| FCs09 | 303.5 | 1.28 | 335.0 | 36.7 | 126.3 | 0.025 | 0.57 | 11.1 | 74.3 | 8.2 |
| **Salinized** |  |  |  |  |  |  |  |  |  |  |
| FCs14 | 2113.4 | 10.60 | 1076.3 | 135.3 | 313.6 | 0.241 | 1.54 | 32.3 | 114.2 | 6.7 |
| FCs10 | 2688.6 | 11.40 | 1462.6 | 213.8 | 535.4 | 0.071 | 1.80 | 41.8 | 137.2 | 6.1 |
| FCs05 | 13824.0 | 51.20 | 6561.9 | 1027.2 | 1116.0 | 0.247 | 6.48 | 114.6 | 396.9 | 5.1 |

By considering all major and trace chemical components (Supplementary Table 1), the chemical composition differed significantly between the two aquifer units (ANOSIM test, p = 0.029). The molar ion ratios Ca/Br, Cl/Ca, Na/Ca, K/Br, Mg/Ca, Cl/K, Br/Sr, Cl/Sr, SO₄/Ca, and Na/K were significantly different between non-salinized and salinized aquifer units (Kruskal-Wallis test, p < 0.05) and accounted for more than 50% of the average group dissimilarity (SIMPER test). Other molar ratios commonly associated with groundwater salinization processes (e.g., seawater intrusion) were less relevant in discriminating the two aquifer units (i.e., Cl/Br, Cl/Na, Cl/HCO_3_).

### 3.2. Effects of historical salinization on microbial community structure

Total cell counts (TCC) ranged from 4.1 to 46.6 × 10^5^ cells/ml, but no significant differences were found between the two sample groups (8.2 ± 3.1 × 10^5^ cells/ml in non-salinized samples and 23.1 ± 20.0 × 10^5^ cells/ml in salinized samples). LNA cells represented the major fraction of total cells (76.6 ± 6.2% of TCC). The occurrence of HNA cells was statistically higher in non-salinized than salinized samples (27.7 ± 2.9% and 17.8 ± 4.5% of TCC, respectively), with a significant shift of the LNA/HNA cell ratio passing from non-salinized to salinized aquifer units (from 2.6 ± 0.4% to 4.9 ± 1.5% of TCC, respectively; Kruskal–Wallis test, p < 0.05).

By considering 71 phyla, 140 classes, 313 orders, and 359 families, we found distinct patterns of microbial community composition between the two aquifer units. The alpha diversity decreased with salinization (from H_phylo_ = 3.5 ± 0.4 to H_phylo_ = 3.0 ± 0.5), and the microbial community composition at the ASV level differed significantly between non-salinized and salinized samples (ANOSIM test, p = 0.023). When restricting the analysis to classes represented in three or more distinct samples and focusing on dominant families (those exceeding 1% relative abundance in at least one sample), we identified 31 dominant families, grouped into five phylogenetic classes (Figure 2). The microbial community of non-salinized samples was significantly enriched in members of the classes Alphaproteobacteria (family Rhizobiaceae), Bacteroidia (family Chitinophagaeae), Clostridia (family Clostridiaceae), Gammaproteobacteria (families Oxalobacteraceae, Moraxellaceae, and Chromobacteraceae), and Methylomirabilia (family Methylomirabiliaceae) (Kruskal–Wallis test, p < 0.05). The microbial community of salinized samples was significantly enriched in members of the classes Campylobacteria (families Sulforimonadaceae, Sulfurovaceae, and Arcobacteraceae) and Gammaproteobacteria (family Nitrincolaceae) (Kruskal–Wallis test, p < 0.05). In the most saline sample (FCs05), Vibrionaceae and Sulfurimonadaceae emerged as the dominant families, collectively comprising the majority of the community, alongside Paracoccaceae and Thalassospiraceae (Figure 2).

**Figure 2.**
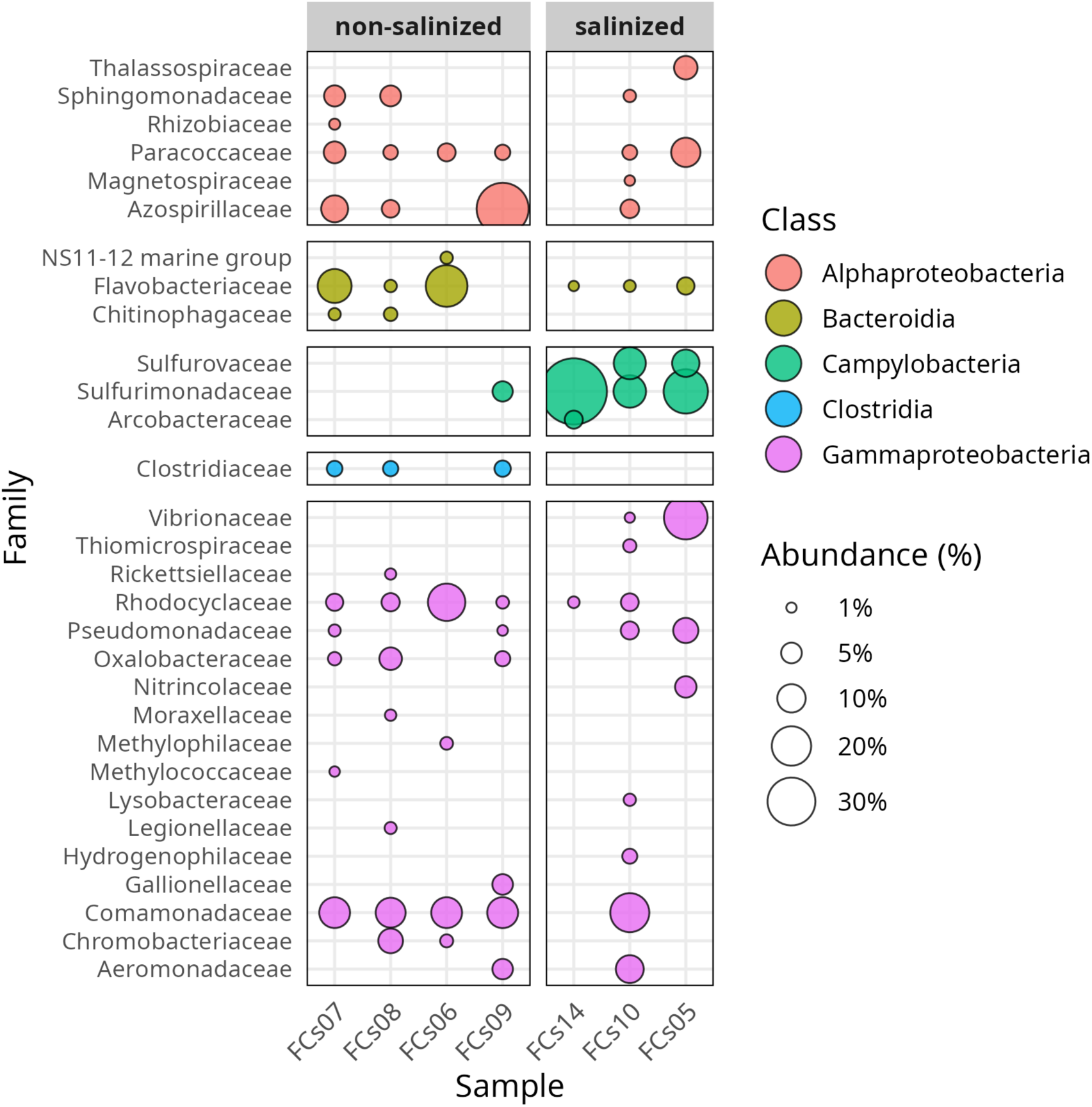
Relative abundance of the dominant phylogenetic bacterial taxa in non-salinized and salinized aquifer units. Samples are ordered left to right by their Cl levels. The classes represented in at least three different samples were included in the visualization, as were families with relative abundances higher than 1% of total reads in at least one sample. The relative abundance values determined the size of the family bubbles, while the corresponding class determined their color.

Based on the LEfSe findings, we identified freshwater-adapted taxa as those significantly enriched in non-salinized samples and halotolerant taxa as those significantly enriched in salinized samples (Figure 3). The freshwater-adapted taxa were relatively more diversified and showed generally low abundance values. The halotolerant taxa were relatively more abundant and almost exclusively affiliated with the phylum Campylobacterota (order Campylobacterales, class Campylobacteria, families Sulfurimonadaceae and Sulfurovaceae). *Sulfurimonas* spp. and *Sulfurovum* spp. were found to be the most abundant genera in salinized samples (Figure 3).

**Figure 3.**
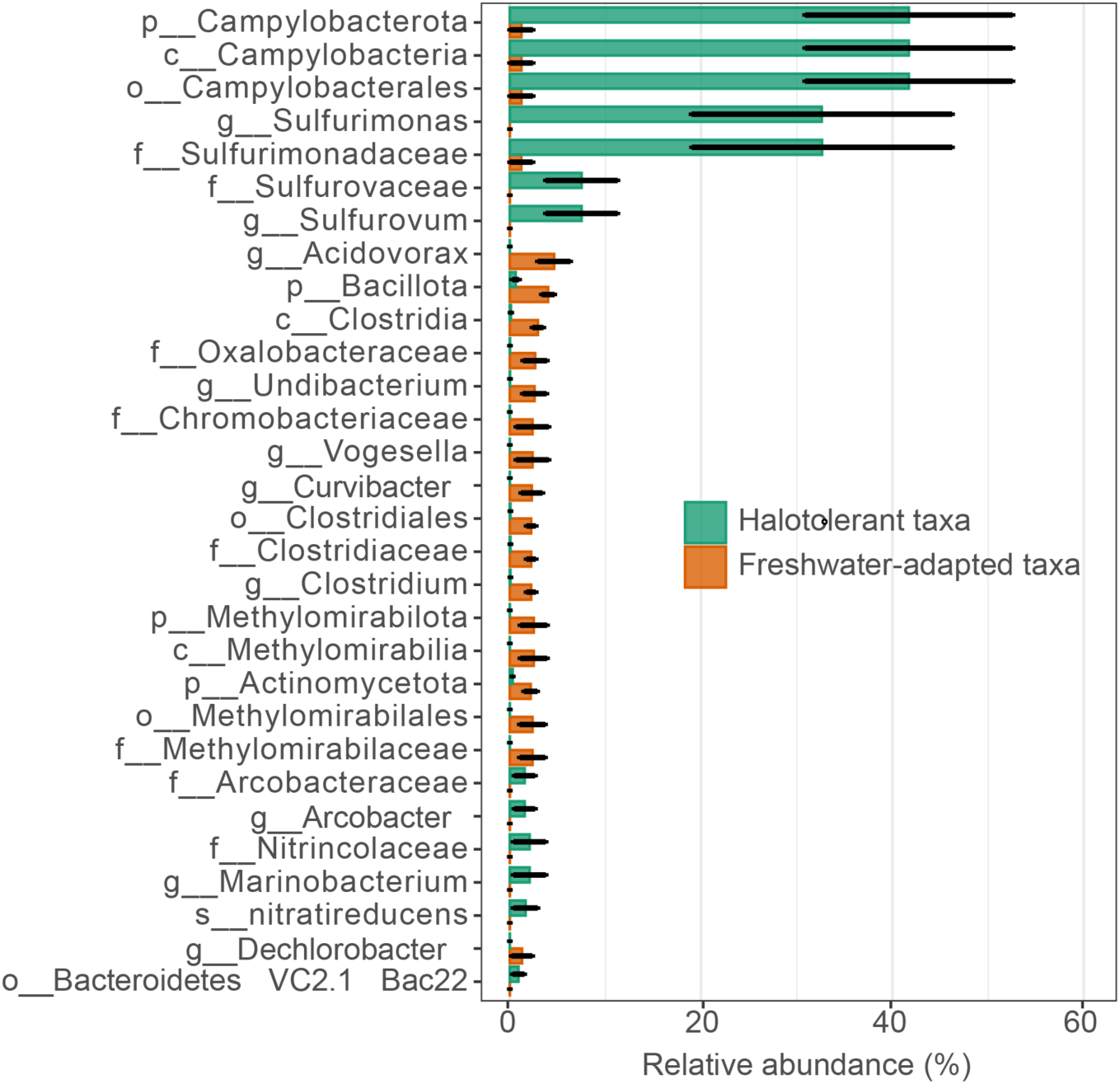
Linear Discriminant Analysis Effect Size (LEfSe) analysis applied to statistically discriminate between freshwater-adapted and halotolerant taxa, respectively enriched in non-salinized and salinized samples. Error bars indicate standard errors.

The cumulative relative abundance at the family level revealed that the four freshwater-adapted families (i.e., Oxalobacteraceae, Methylomirabilaceae, Clostridiaceae, and Chromobacteriaceae) comprised 9.6 ± 4.5% of the total reads in non-salinized samples, which declined significantly in salinized samples (0.09 ± 0.06% of total reads).

In contrast, members of the four halotolerant families (i.e., Sulfurimonadaceae, Sulfurovaceae, Arcobacteraceae, and Nitrincolaceae) predominated in salinized samples (43.8 ± 17.7% of total reads), whereas they constituted 1.2 ± 2.4% of total reads in non-salinized samples. The cumulative abundances of freshwater-adapted and halotolerant families were inversely correlated (r_s_ = -0.93, p = 0.007).

### 3.3. Effects of historical salinization on microbial metabolic potential and functional profiles

The FAPROTAX functions related to the carbon, nitrogen, and sulfur cycles did not differ between the two aquifer units, as indicated by both univariate and multivariate analyses (Kruskal-Wallis and ANOSIM tests, p > 0.05). The annotated carbon-related functions were prevalent, with elevated values for chemoheterotrophy, fermentation, and hydrocarbon degradation. Nitrogen- and sulfur-related functions primarily involved reduction and respiration processes, specifically denitrification and sulfate respiration (Figure 4).

**Figure 4.**
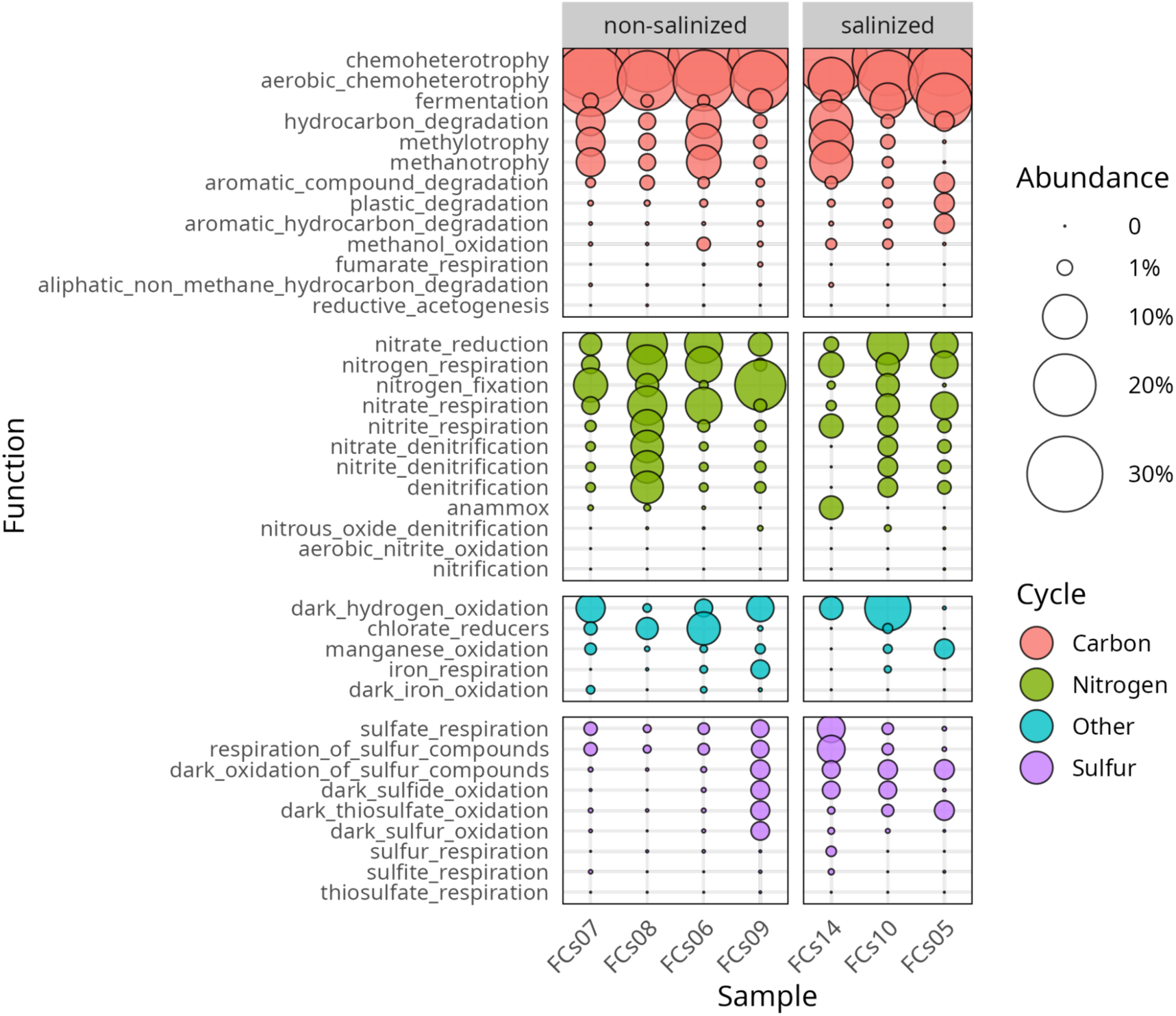
FAPROTAX-annotated functions were categorized into groups associated with primary metabolic processes, mainly involved in carbon, nitrogen, and sulfur cycles. Samples are grouped in non-salinized and salinized aquifer units and ordered left to right by their Cl levels. The bubble size is scaled according to the relative abundance of total annotated functions.

The Biolog assays validated the microbial metabolic capacity to process organic matter and carbon-based compounds, whether naturally occurring (MT2 tests) or artificially introduced (EcoPlates tests), via chemoheterotrophic mechanisms. The heterotrophic respiration was significantly lower in non-salinized samples compared to salinized ones (0.06 ± 0.03 OD_590nm_ and 0.46 ± 0.10 OD_590nm_, respectively; Kruskal–Wallis test, p < 0.05). The same pattern was observed for the AWCD (0.04 ± 0.01 OD_590nm_ and 1.85 ± 0.67 OD_590nm_, respectively; Kruskal–Wallis test, p < 0.05). Non-salinized samples exhibited a more extensive utilization of carbon sources, demonstrating significantly elevated functional diversity values in comparison to salinized samples (H_funct_ = 1.9 ± 0.2 and H_funct_ = 1.6 ± 0.1 respectively; Kruskal–Wallis test, p < 0.05). Notably, the high salinity level significantly promoted the degradation of polymers, with absorbance values increasing from 0.02 ± 0.01 OD_590nm_ in non-salinized samples to 0.13 ± 0.01 OD_590nm_ in salinized ones (Figure 5).

**Figure 5.**
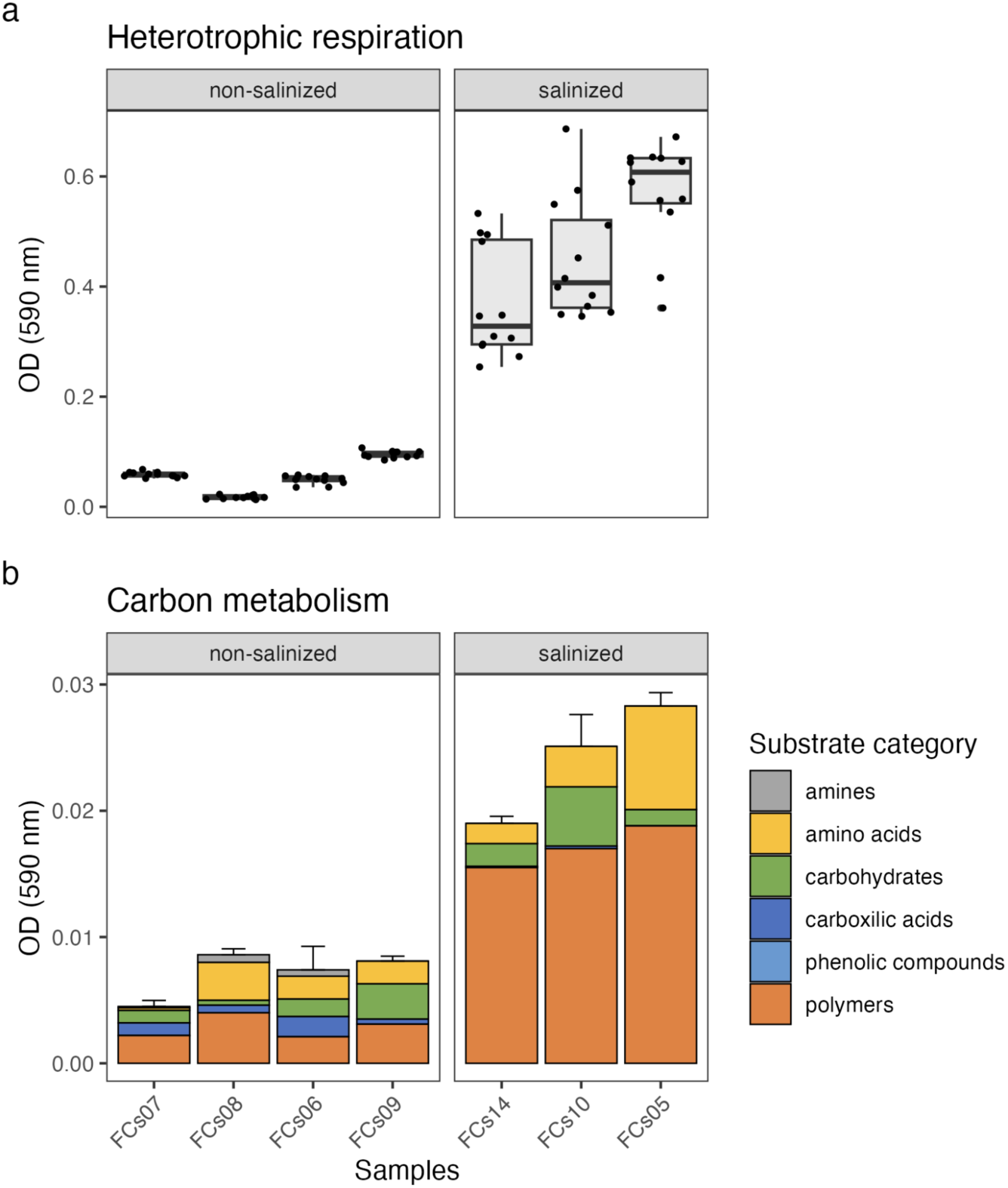
Microbial functional profiles in non-salinized and salinized aquifer units. Samples are ordered left to right by their Cl levels. Heterotrophic respiration (a) was assessed using the Biolog MT2 assay. The carbon metabolism (b) was evaluated based on the degradation of Biolog EcoPlates substrates, grouped according to the substrate category.

### 3.4. Multivariate statistics for data integration

The NMDS ordination plot was applied to graphically visualize the effects of historical salinization on groundwater chemical composition and microbial community profiles (stress value < 0.01). The primary separation in the ordination space took place along the first NMDS axis, which significantly differentiated between the two aquifer units (ANOSIM test, p = 0.028). Vectors representing chemical variables (i.e., relevant molar ratios) and microbial community descriptors (i.e., relevant structural and functional characteristics) were differently oriented among non-salinized and salinized samples, showing significant associations with the local environment. The clear separation of non-salinized and salinized aquifer units along the groundwater salinization gradient revealed coordinated responses, with divergent chemical and microbial vectors tracking geochemical conditions in microbial community profiles (Figure 6).

**Figure 6.**
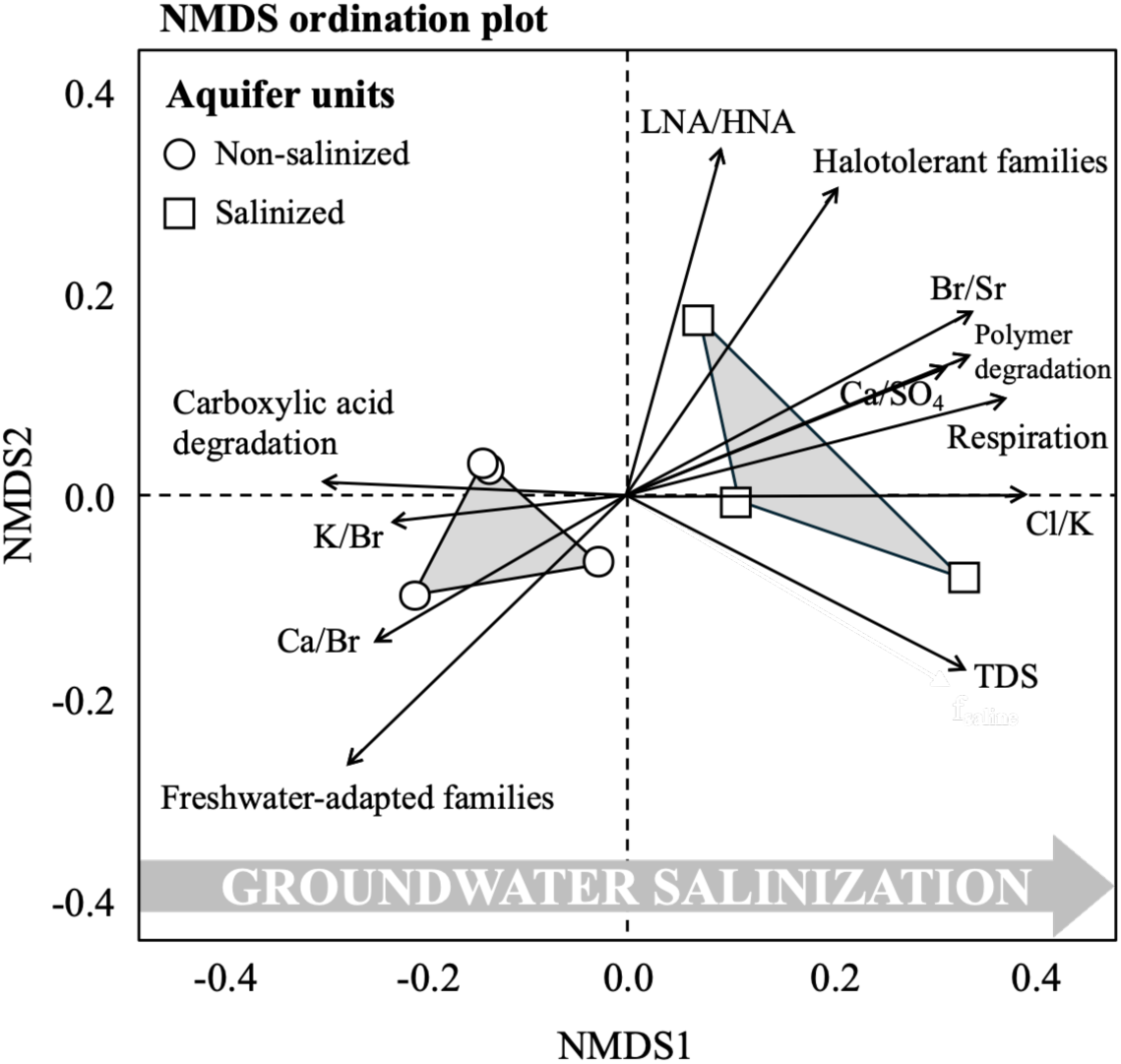
NMDS ordination plot illustrating the impact of historical groundwater salinization on chemical and microbial community profiling. Samples were grouped by non-salinized and salinized aquifer units. The environmental vectors represent the key parameters for chemical composition (Ca/SO_4_, K/Br, Ca/Br, Br/Sr, Cl/K, TDS), microbial community structure (LNA/HNA cell ratio, the cumulative abundance of freshwater-adapted and halotolerant families), and degradative metabolic potential (heterotrophic respiration, carboxylic acid and polymer degradation capacity) that most effectively differentiated the two aquifer units.

## 4. Discussion

The patterns of hydrogeochemical data and microbial community profiles demonstrated that historical salinization from ancient saltworks continues to structure the present-day aquifer ecosystem through coupled geochemical and biological pathways. The ecological legacy of ancient saltworks can extend far beyond the salt accumulation, creating persistent ecosystems with fundamentally altered geochemistry, microbial community composition, metabolic pathways, and functional capacities (Jakab et al., 2020; Kaushal et al., 2023). The two-millennia timespan since the cessation of Roman salt production activities demonstrated the extraordinary temporal persistence of anthropogenic impacts on groundwater ecosystems, as groundwater systems show far slower dynamics than surface environments due to limited connectivity, slow flow rates, and the persistence of salt deposits that continue to dissolve over geological timescales (Robinson et al., 2017; Erostate et al., 2018; Seibert et al., 2024).

### 4.1. Historical salinization signatures in groundwater chemistry

The increase of salinity with distance from the sea rather than proximity to it provided evidence that the observed salinization was not driven by contemporary seawater intrusion but rather by historical salt sources. The elevated ionic strength and enriched concentrations of halogen (i.e., Cl, Br) and alkaline (i.e., Na, K) ions in the salinized aquifer unit indicated the persistent geochemical signature of historical saltwork operations that previously prevailed in the target coastal region (Jasechko et al., 2017; Xu et al., 2021).

Distinctive ionic ratio patterns were recognized as fingerprints of historical salinization processes rather than active seawater intrusion (Argamasilla et al., 2017; Selvakumar et al., 2022). In this study, Cl/Na molar ratio was significantly different in the two aquifer units, with the ratio in saline samples close to that of seawater, while Cl/Br molar ratio was significantly different from the seawater one only in the most fresh samples, supporting a recent source of saline water. This is in line with recent findings demonstrating that paleointrusions and relic saline deposits can exhibit hydrochemical signatures that deviate substantially from typical marine ratios due to diagenetic processes, selective ion retention in sediments, and differential ion mobility through aquitards over geological timescales (Renau-Pruñonosa et al., 2025; Wen et al., 2025). The enrichment of specific elemental ratios in our salinized samples (i.e., Br/Sr, Cl/K, and SO₄/Ca) suggested complex geochemical processes associated with the dissolution and mobilization of historical salt deposits (Wu and Wang, 2014; Ongetta et al., 2022). Ancient Roman saltworks typically involved the concentration of seawater in shallow evaporation ponds, where progressive evaporation led to sequential precipitation of different mineral phases (e.g., carbonates, gypsum, and halite), creating stratified salt deposits with distinctive chemical compositions (Vicari et al., 2022). In salinized samples, the elevated SO₄/Ca ratios indicated preferential dissolution processes, while the modified Mg/Ca and K/Br ratios reflected the influence of late-stage evaporite minerals that form during hypersaline conditions. Recent studies indicated that evaporite minerals and anthropogenic saline deposits can drive groundwater chemistry for centuries after their cultural and economic end points (Ahmed et al., 2013; Alessandrino et al., 2023; Abidi et al., 2024).

### 4.2. Microbial community restructuring in response to historical salinization

The microbial community analysis revealed significant structural reorganization between non-salinized and salinized aquifer units, despite the maintenance of comparable total cell counts across the salinity gradient. In saline conditions, the significant increase of the LNA/HNA cell ratio, alongside the high microbial metabolic potential, indicated that historical salinization was likely to alter the microbial community structure, possibly reflecting a phylogenetic shift with the selection of microbial taxa inherently characterized by a low per-cell content of nucleic acids. In previous studies, the salinity fluctuations were found to regulate the ecological functions and the interaction dynamics between LNA and HNA cells in surface waters (Hu et al., 2024, 2025). Notably, LNA cells were reported as small-sized microorganisms with slow metabolic functions and diverse survival strategies for thriving in groundwater-dependent ecosystems (Di Lorenzo et al., 2025).

Relevant structural changes occurred in the phylogenetic community composition as also assessed by 16S rRNA gene amplicon sequencing. The taxonomic analysis identified distinct freshwater-adapted and halotolerant assemblages that characterized the two aquifer units. The non-salinized samples harbored a relatively diverse suite of freshwater-adapted families, including Oxalobacteraceae, Methylomirabilaceae, Clostridiaceae, and Chromobacteriaceae, spanning multiple proteobacterial classes as well as Clostridia and the methane-oxidizing Methylomirabilia. These taxa were commonly associated with freshwater sediments and groundwater systems, where they were recognized for their involvement in carbon metabolic processes, including organic matter degradation, methanotrophy, and fermentation (Percak-Dennett and Roden, 2014; Barnett et al., 2021). In contrast, the salinized samples were dominated by a taxonomically narrow but highly abundant assemblage of halotolerant taxa almost exclusively affiliated with the phylum Campylobacterota, particularly the families Sulfurimonadaceae and Sulfurovaceae. Within these families, the genera *Sulfurimonas* and *Sulfurovum* emerged as the most abundant components of the salinized aquifer microbiome. Members of these bacterial taxa were recognized as halotolerant sulfur-processing bacteria that typically occupy ecological niches at oxic-anoxic interfaces (Hamilton et al., 2015; Alamoudi et al., 2025). The inverse correlation between the occurrence of freshwater-adapted and halotolerant families was likely to indicate the competitive displacement of freshwater taxa by salt-tolerant specialists as salinity increased. This pattern reflects fundamental osmoregulatory constraints on microbial physiology, since freshwater-adapted microorganisms may lack the genetic suite necessary to synthesize or accumulate the compatible solutes required to maintain osmotic balance under elevated salinity (Wang et al., 2023; West et al., 2025).

Previous studies on coastal aquifers subject to seawater intrusion showed that members of the class Gammaproteobacteria (i.e., family Alteromonadaceae) and Alphaproteobacteria (i.e. family Rhodobacteraceae) became increasingly abundant at rising salinity levels (Chen et al., 2019; Zhi et al., 2024). Here, the enrichment of halotolerant microorganisms occurred mostly within the class Campylobacteria rather than gammaproteobacterial lineages, which were found to prevail in systems where organic carbon serves as the primary energy source (Risley et al., 2022). This observation underscores the importance of considering both salinity levels and associated geochemical conditions when predicting microbial community responses to groundwater salinization (Houghton et al., 2023).

### 4.3. Functional resilience and metabolic specialization under saline conditions

The functional characterization revealed a complex relationship between microbial community structure and metabolic potential across the salinity gradient. While the taxonomic composition underwent consistent restructuring, the FAPROTAX-annotated functions associated with carbon, nitrogen, and sulfur cycling showed no significant differences between non-salinized and salinized aquifer units. In line with literature findings (Li et al., 2024), carbon-related functions (i.e., chemoheterotrophy, fermentation, and hydrocarbon degradation) dominated across both aquifer units, with a broad occurrence of nitrogen- and sulfur-related functions (i.e., denitrification and sulfate respiration). This apparent functional stability despite taxonomic shifts suggested a degree of functional redundancy in the groundwater microbial community. Phylogenetically distinct taxa were likely to maintain similar metabolic capabilities, thereby preserving ecosystem-level biogeochemical processes even at increasing salinity levels (Yang et al., 2022; Zhi et al., 2024; Selak et al., 2025).

However, the gene-centric FAPROTAX analysis failed to capture the consistent functional differences revealed by the Biolog assays. When measuring the potential metabolic activity under controlled laboratory conditions, heterotrophic respiration and average well color development (AWCD) were significantly lower in non-salinized samples compared to salinized ones. Our results indicated substantially increasing metabolic activity and carbon substrate utilization capacity with increasing salinity levels. The elevated metabolic activity was likely to reflect the adaptations of halotolerant communities to carbon limitation in saline environments, as microorganisms adapted to high-salinity conditions can maintain high expression levels of catabolic enzymes and transport systems to maximize carbon and energy acquisition efficiency under osmotically stressful conditions where osmoregulation imposes substantial energetic costs (Gunde-Cimerman et al., 2018; Xamxidin et al., 2025).

The functional diversity analysis revealed that non-salinized samples exhibited significantly higher H_funt_ values compared to salinized samples, indicating a broader range of pathways for utilizing carbon substrates in freshwater-adapted communities. The microbial community in salinized samples showed higher polymer degradation capacity, with absorbance values increasing more than sixfold compared to non-salinized samples. Polymers are complex, high-molecular-weight organic compounds that require extracellular enzymatic depolymerization before cellular uptake and metabolism (Hino et al., 2023). The elevated polymer degradation capacity in salinized samples suggested enrichment of taxa producing extracellular hydrolytic enzymes, which may be ecologically significant in groundwater environments where organic matter can occur in particulate or polymeric forms (Zoppini et al., 2020; Wang et al., 2024).

These findings were likely to address a critical knowledge gap regarding whether halotolerant species can fully compensate for the ecological roles of taxa that decline under elevated salinity (Izabel-Shen, 2021; Nelson et al., 2024). In this study, the microbial community in salinized conditions demonstrated reduced metabolic versatility but enhanced capacity for specific metabolic processes that may be adaptive under saline conditions. This functional reorganization implied that while basic biogeochemical cycling may persist, the specific pathways and ecological outcomes may differ at contrasting salinity levels (Zhi et al., 2024). This transition represented a fundamental shift in ecosystem functioning, exploiting a subsurface environment characterized by aphotic, anoxic, and high salinity conditions. Such polyextreme ecosystems, where microbes form the base of the food web independently of photosynthetically derived organic matter, are relatively rare but well-documented in inland aquatic ecosystems, such as bottom waters of saline meromictic lakes (Fazi et al., 2021), sulfide-rich caves (Jones et al., 2014), blue holes (Björnerås et al., 2020), deep saline aquifers (Payler et al., 2019), and halite deposits (Dindhoria et al., 2023).

### 4.4. Chemical–microbial coupling driven by historical salinization and implications for groundwater management and ecosystem services

The coupled chemical-microbial signatures suggested that historical salinization created alternative ecosystem states within different units of the same aquifer, owing to freshwater recharge in the non-salinized unit and salt dissolution processes in the salinized unit. A critical challenge highlighted by our results is the difficulty of distinguishing historical salinization from active seawater intrusion using conventional hydrochemical indicators.

Owing to a limited discriminatory power, aquifer salinization may be misattributed to current groundwater over-extraction and seawater intrusion when the actual cause might be the dissolution of buried evaporite deposits from historical activities and residual brines (Chandrajith et al., 2016; Alessandrino et al., 2023). This misdiagnosis can have important effects on how to manage the local groundwater resources, since strategies that aim to stop seawater intrusion by lowering pumping or injection barriers would not work against salinization from distributed evaporite sources (Mastrocicco and Colombani, 2021; Narvaez-Montoya et al., 2023).

Our results demonstrated that a comprehensive groundwater characterization, including chemical and microbial community descriptors, can be essential for accurate source attribution and appropriate management response selection. For instance, the site FCs09 showed a moderate saline water mixing fraction (f_saline_ = 0.8%, TDS = 1448 mg/l), alongside a relatively high content in alkalis (Na/Cl molar ratio = 1.7). This suggested saline groundwater refreshening, with release of Na previously adsorbed to sediment particles in that particular location of the study area (Jiao and Post, 2019). However, the microbial community profiles at FCs09 more closely resembled those of fresh groundwater samples. This discrepancy suggested that aquatic microorganisms could respond more rapidly to ongoing refreshening processes, while hydrochemical signatures remain more influenced by historical salinization effects (Zheng et al., 2019; Bourhane et al., 2023; Chiriac et al., 2023). Moreover, the presence of halotolerant taxa, such as the families Sulfurimonadaceae and Sulfurovaceae, could serve as biological indicators of salinization, potentially offering further insights to complement chemical assessments (Rossmassler et al., 2016). Although these observations should be approached with caution given the limited number of analyzed samples, the underestimated ecological responsiveness could make microbial indicators especially useful for identifying underrated changes in aquifer conditions (Zheng et al., 2019; Xiu et al., 2020).

Despite the insights provided by this study, investigation across diverse geological settings is needed to assess the generality of our findings, as aquifer lithology, mineralogy, and sediment characteristics likely influence both the geochemical impacts of salinization and the microbial community responses. The overall understanding of community assembly dynamics may enhance predictions regarding microbial community responses to current salinization processes, including ongoing seawater intrusion driven by climate change and groundwater overexploitation (Ghirardelli et al., 2025).

## 5. Conclusions

This study demonstrates that ancient saltworks can create a lasting ecological legacy that continues to shape coastal aquifer ecosystems, fundamentally restructuring microbial communities and altering ecosystem functions in ways that affect present-day water resources and ecosystem services. Our findings point out the importance of historical awareness in groundwater management and the value of integrated chemical-biological monitoring approaches to recognizing the potential for anthropogenic impacts to persist across geological timescales.

## Data Availability

The sequencing data from this study have been deposited in the European Nucleotide Archive (ENA) and can be accessed under BioProject PRJEB98918. All data and codes used for statistical and bioinformatic analyses can be retrieved in the following GitHub Repository (https://github.com/MicroEcoLab/Fiumicino-aquifer).

## Acknowledgments

This study was funded by a joint local project between CNR-IRSA and Aeroporti di Roma (AdR) aimed at defining the natural geochemical background levels of groundwater at the area surrounding the Fiumicino airport. The experimental work was also supported by the project “National Biodiversity Future Center - NBFC” under the National Recovery and Resilience Plan (NRRP). We thank the Fiumicino airport personnel for their kind support and for granting access to the sampling area.

## Supplementary Material

**Supplementary Table 1.** Field data and physical-chemical descriptors. Sampling sites are ordered by the increasing salinity level (a). Major chemical components (b), trace elements (c), and molar ion ratios (d) measured in non-salinized and salinized groundwater samples. Parameters are ordered left to right by their relative contribution to average group dissimilarity (SIMPER test). Asterisks indicate significant differences between median values between the two aquifer units (Kruskal–Wallis test, p < 0.05). The complete dataset used for statistical analyses can be retrieved in the GitHub Repository - https://github.com/MicroEcoLab/Fiumicino-aquifer.

| (a) | Lat.<br>UTM ED50 | Long.<br>UTM ED50 | Elevation<br>m a.s.l. | T<br>°C | pH<br>- | ORP<br>mV | *EC<br>μS/cm |
| --- | --- | --- | --- | --- | --- | --- | --- |
| **Non-salinized** |  |  |  |  |  |  |  |
| FCs07 | 272005 | 4631509 | 0.14 | 18.6 | 6.8 | 96.0 | 640 |
| FCs08 | 271050 | 4631279 | 1.32 | 19.1 | 9.7 | 243.0 | 516 |
| FCs06 | 270508 | 4631198 | 1.0 | 21.0 | 7.5 | -186.0 | 669 |
| FCs09 | 272670 | 4631728 | 0.9 | 19.9 | 7.1 | -201.0 | 2130 |
| **Salinized** |  |  |  |  |  |  |  |
| FCs14 | 274083 | 4631230 | 0.59 | 24.9 | 7.0 | 161.0 | 6778 |
| FCs10 | 273585 | 4632166 | 0.21 | 19.9 | 6.8 | -192.6 | 8560 |
| FCs05 | 273933 | 4632657 | 0.71 | 20.7 | 7.1 | -190.0 | 32500 |

| (b) | *Cl<br>mg/l | *Na<br>mg/l | *SO<sub>4</sub><br>mg/l | HCO<sub>3</sub><br>mg/l | *Mg<br>mg/l | *Ca<br>mg/l | *K<br>mg/l | *Br<br>mg/l | NO<sub>3</sub><br>mg/l | *Si<br>mg/l | NH<sub>4</sub><br>mg/l | O<sub>2</sub><br>mg/l | F<br>mg/l | PO<sub>4</sub><br>mg/l |
| --- | --- | --- | --- | --- | --- | --- | --- | --- | --- | --- | --- | --- | --- | --- |
| **Non-salinized** |  |  |  |  |  |  |  |  |  |  |  |  |  |  |
| FCs07 | 13.6 | 14.3 | 13.0 | 338.2 | 8.4 | 89.7 | 12.3 | 0.04 | 9.8 | 11.0 | 0.1 | 0.2 | 0.2 | 0.2 |
| FCs08 | 16.6 | 10.6 | 7.4 | 249.9 | 11.1 | 59.4 | 4.0 | 0.03 | 19.8 | 9.0 | 0.0 | 1.1 | 0.1 | 0.3 |
| FCs06 | 31.1 | 27.8 | 8.1 | 335.4 | 13.8 | 75.8 | 7.6 | 0.23 | 0.6 | 10.2 | 0.2 | 0.7 | 0.3 | 0.1 |
| FCs09 | 303.5 | 335.0 | 126.3 | 766.6 | 36.7 | 74.3 | 11.1 | 1.28 | 1.1 | 8.2 | 0.8 | 0.7 | 1.2 | 0.1 |
| **Salinized** |  |  |  |  |  |  |  |  |  |  |  |  |  |  |
| FCs14 | 2113.4 | 1076.3 | 313.6 | 384.3 | 135.3 | 114.2 | 32.3 | 10.60 | 9.0 | 6.7 | 0.7 | 1.5 | 0.4 | 0.9 |
| FCs10 | 2688.6 | 1462.6 | 535.4 | 871.1 | 213.8 | 137.2 | 41.8 | 11.40 | 3.2 | 6.1 | 1.4 | 0.9 | 0.9 | 0.1 |
| FCs05 | 13824.0 | 6561.9 | 1116.0 | 1348.8 | 1027.2 | 396.9 | 114.6 | 51.20 | 9.3 | 5.1 | 0.5 | 1.9 | 0.1 | 0.1 |

| (c) | *Sr<br>μg/l | Fe<br>μg/l | B<br>μg/l | Mn<br>μg/l | *Al<br>μg/l | Ba<br>μg/l | Li<br>μg/l | V<br>μg/l | Se<br>μg/l |
| --- | --- | --- | --- | --- | --- | --- | --- | --- | --- |
| **Non-salinized** |  |  |  |  |  |  |  |  |  |
| FCs07 | 283.1 | 62.5 | 75.0 | 304.3 | 6.2 | 39.5 | 4.1 | 4.5 | 3.7 |
| FCs08 | 260.5 | 18.8 | 75.0 | 158.7 | 17.5 | 19.0 | 1.0 | 2.7 | 0.5 |
| FCs06 | 217.1 | 699.5 | 75.0 | 496.2 | 12.8 | 25.4 | 4.1 | 1.0 | 0.5 |
| FCs09 | 568.0 | 2512.8 | 940.7 | 200.1 | 24.8 | 165.4 | 26.2 | 2.2 | 4.3 |
| **Salinized** |  |  |  |  |  |  |  |  |  |
| FCs14 | 1535.8 | 337.1 | 742.1 | 279.4 | 240.6 | 83.9 | 21.3 | 11.1 | 0.5 |
| FCs10 | 1804.4 | 2595.3 | 1404.6 | 774.2 | 70.7 | 176.8 | 52.4 | 10.0 | 12.8 |
| FCs05 | 6483.4 | 638.1 | 1807.5 | 1130.3 | 246.6 | 275.0 | 129.4 | 1.0 | 11.6 |

| (d) | *Ca/Br | *Cl/Ca | *Na/Ca | *K/Br | *Mg/Ca | *Cl/K | *Br/Sr | *Cl/Sr | *SO<sub>4</sub>/Ca |
| --- | --- | --- | --- | --- | --- | --- | --- | --- | --- |
| **Non-salinized** |  |  |  |  |  |  |  |  |  |
| FCs07 | 8938 | 0.09 | 0.14 | 628.3 | 0.15 | 1.22 | 0.04 | 30.0 | 0.06 |
| FCs08 | 7892 | 0.16 | 0.16 | 272.4 | 0.31 | 4.58 | 0.03 | 39.5 | 0.05 |
| FCs06 | 1314 | 0.23 | 0.32 | 67.5 | 0.30 | 4.51 | 0.29 | 87.4 | 0.04 |
| FCs09 | 231 | 2.31 | 3.93 | 17.7 | 0.81 | 30.16 | 0.62 | 329.2 | 0.71 |
| **Salinized** |  |  |  |  |  |  |  |  |  |
| FCs14 | 43 | 10.47 | 8.21 | 6.2 | 1.95 | 72.18 | 1.89 | 848.5 | 1.15 |
| FCs10 | 48 | 11.08 | 9.29 | 7.5 | 25.69 | 70.96 | 1.74 | 923.6 | 1.63 |
| FCs05 | 31 | 19.70 | 14.41 | 4.6 | 4.27 | 133.1 | 2.17 | 1319.1 | 1.17 |

## References

1. Abidi, J. H., Elzain, H. E., Sabarathinam, C., El Fehri, R. M., Farhat, B., Ben Mammou, A., et al. (2024). Integrated approach to understand the multiple natural and anthropogenic stresses on intensively irrigated coastal aquifer in the Mediterranean region. Environmental Research 252, 118757. doi: 10.1016/j.envres.2024.118757

2. Ahmed, M. A., Abdel Samie, S. G., and Badawy, H. A. (2013). Factors controlling mechanisms of groundwater salinization and hydrogeochemical processes in the Quaternary aquifer of the Eastern Nile Delta, Egypt. Environ Earth Sci 68, 369–394. doi: 10.1007/s12665-012-1744-6

3. Alamoudi, R., Barozzi, A., Michoud, G., Van Goethem, M. W., Odobel, C., Chen, Y., et al. (2025). Metabolic redundancy and specialisation of novel sulfide-oxidizing Sulfurimonas and Sulfurovum along the brine-seawater interface of the Kebrit Deep. Environmental Microbiome 20, 19. doi: 10.1186/s40793-025-00669-7

4. Alessandrino, L., Gaiolini, M., Cellone, F. A., Colombani, N., Mastrocicco, M., Cosma, M., et al. (2023). Salinity origin in the coastal aquifer of the Southern Venice lowland. Science of The Total Environment 905, 167058. doi: 10.1016/j.scitotenv.2023.167058

5. Amalfitano, S., Del Bon, A., Zoppini, A., Ghergo, S., Fazi, S., Parrone, D., et al. (2014). Groundwater geochemistry and microbial community structure in the aquifer transition from volcanic to alluvial areas. Water Research 65, 384–394. doi: 10.1016/j.watres.2014.08.004

6. Amalfitano, S., Fazi, S., Ejarque, E., Freixa, A., Romaní, A. M., and Butturini, A. (2018). Deconvolution model to resolve cytometric microbial community patterns in flowing waters. Cytometry Pt A 93, 194–200. doi: 10.1002/cyto.a.23304

7. Antonites, A. (2013). Archaeological salt production at the Baleni spring, northeastern South Africa. South African Archaeological Bulletin 68, 105–118.

8. Appelo, C. A. J., and Postma, D. (2004). Geochemistry, Groundwater and Pollution., 0 Edn, eds. C. A. J. Appelo and D. Postma. CRC Press. doi: 10.1201/9781439833544

9. Argamasilla, M., Barberá, J. A., and Andreo, B. (2017). Factors controlling groundwater salinization and hydrogeochemical processes in coastal aquifers from southern Spain. Science of The Total Environment 580, 50–68. doi: 10.1016/j.scitotenv.2016.11.173

10. Barnett, S. E., Youngblut, N. D., Koechli, C. N., and Buckley, D. H. (2021). Multisubstrate DNA stable isotope probing reveals guild structure of bacteria that mediate soil carbon cycling. Proc. Natl. Acad. Sci. U.S.A. 118, e2115292118. doi: 10.1073/pnas.2115292118

11. Bierkens, M. F. P., and Wada, Y. (2019). Non-renewable groundwater use and groundwater depletion: a review. Environ. Res. Lett. 14, 063002. doi: 10.1088/1748-9326/ab1a5f

12. Björnerås, C., Škerlep, M., Gollnisch, R., Herzog, S. D., Ekelund Ugge, G., Hegg, A., et al. (2020). Inland blue holes of The Bahamas – chemistry and biology in a unique aquatic environment. Fundamental and Applied Limnology 194, 95–106. doi: 10.1127/fal/2020/1330

13. Bourhane, Z., Cagnon, C., Castañeda, C., Rodríguez-Ochoa, R., Álvaro-Fuentes, J., Cravo-Laureau, C., et al. (2023). Vertical organization of microbial communities in Salineta hypersaline wetland, Spain. Front. Microbiol. 14, 869907. doi: 10.3389/fmicb.2023.869907

14. Callahan, B. J., McMurdie, P. J., Rosen, M. J., Han, A. W., Johnson, A. J. A., and Holmes, S. P. (2016). DADA2: High-resolution sample inference from Illumina amplicon data. Nat Methods 13, 581–583. doi: 10.1038/nmeth.3869

15. Chandrajith, R., Diyabalanage, S., Premathilake, K. M., Hanke, C., Van Geldern, R., and Barth, J. A. C. (2016). Controls of evaporative irrigation return flows in comparison to seawater intrusion in coastal karstic aquifers in northern Sri Lanka: Evidence from solutes and stable isotopes. Science of The Total Environment 548-549, 421–428. doi: 10.1016/j.scitotenv.2016.01.050

16. Chen, L., Hu, B. X., Dai, H., Zhang, X., Xia, C.-A., and Zhang, J. (2019). Characterizing microbial diversity and community composition of groundwater in a salt-freshwater transition zone. Science of The Total Environment 678, 574–584. doi: 10.1016/j.scitotenv.2019.05.017

17. Chen, L., Ma, M., Li, X., Yu, K., Zhi, C., Cheng, L., et al. (2024). Effects of Fresh Groundwater and Seawater Mixing Proportions and Salt-Freshwater Displacement on Coastal Aquifer Microbial Communities. Water 16, 2078. doi: 10.3390/w16152078

18. Chiriac, M., Haber, M., and Salcher, M. M. (2023). Adaptive genetic traits in pelagic freshwater microbes. Environmental Microbiology 25, 606–641. doi: 10.1111/1462-2920.16313

19. Corso, D., Melita, M., Massaccesi, N., Quero, G. M., Basili, M., Di Cesare, A., et al. (2025). Constructed wetlands for aquaculture wastewater treatment: insights on the structural and functional shifts of the aquatic microbial community. doi: 10.1101/2025.09.02.673640

20. Cui, Z., Chen, G., Shan, X., Zhang, H., Zhou, Q., Fu, T., et al. (2025). Controls of paleosedimentary environments and anthropogenic activities on coastal groundwater salinization: A case study of Laizhou Bay, China. Marine Geology 487, 107594. doi: 10.1016/j.margeo.2025.107594

21. Dang, X., Gao, M., Wen, Z., Hou, G., Jakada, H., Ayejoto, D., et al. (2022). Saline groundwater evolution in the Luanhe River delta (China) during the Holocene: hydrochemical, isotopic, and sedimentary evidence. Hydrol. Earth Syst. Sci. 26, 1341–1356. doi: 10.5194/hess-26-1341-2022

22. Di Lorenzo, T., Amalfitano, S., Galassi, D. M. P., Melita, M., Zoppini, A., Parrone, D., et al. (2025). Geochemical and microbial factors driving crustacean assemblages in adjacent aquifer units within the same aquifer. Biogeosciences 22, 1237–1256. doi: 10.5194/bg-22-1237-2025

23. Dindhoria, K., Jain, R., Kumar, R., Bhargava, B., Kumar, R., and Kumar, S. (2023). Microbial community structure analysis of hypersaline niches and elucidation of their role in the biogeochemical cycling of nitrogen, sulphur and methane. Ecological Informatics 75, 102023. doi: 10.1016/j.ecoinf.2023.102023

24. Erostate, M., Huneau, F., Garel, E., Lehmann, M. F., Kuhn, T., Aquilina, L., et al. (2018). Delayed nitrate dispersion within a coastal aquifer provides constraints on land-use evolution and nitrate contamination in the past. Science of The Total Environment 644, 928–940. doi: 10.1016/j.scitotenv.2018.06.375

25. Fazi, S., Amalfitano, S., Venturi, S., Pacini, N., Vazquez, E., Olaka, L. A., et al. (2021). High concentrations of dissolved biogenic methane associated with cyanobacterial blooms in East African lake surface water. Commun Biol 4, 845. doi: 10.1038/s42003-021-02365-x

26. Flad, R. K. (2007). Rethinking the Context of Production through an Archaeological Study of Ancient Salt Production in the Sichuan Basin, China. Archaeological Papers of the American 17, 108–128. doi: 10.1525/ap3a.2007.17.1.108

27. Foster, S., Pulido-Bosch, A., Vallejos, Á., Molina, L., Llop, A., and MacDonald, A. M. (2018). Impact of irrigated agriculture on groundwater-recharge salinity: a major sustainability concern in semi-arid regions. Hydrogeol J 26, 2781–2791. doi: 10.1007/s10040-018-1830-2

28. Funiciello, R., and Giordano, G. (2008). Note illustrative della Carta Geologica d’Italia alla scala 1:50.000, Foglio 347 ROMA. Available at: https://www.isprambiente.gov.it/Media/carg/note_illustrative/374_Roma.pdf

29. Gasol, J. M., and Morán, X. A. G. (2015). “Flow Cytometric Determination of Microbial Abundances and Its Use to Obtain Indices of Community Structure and Relative Activity,” in Hydrocarbon and Lipid Microbiology Protocols, eds. T. J. McGenity, K. N. Timmis, and B. Nogales (Berlin, Heidelberg: Springer Berlin Heidelberg), 159–187. doi: 10.1007/8623_2015_139

30. Ghirardelli, A., Straffelini, E., Park, E., D’Agostino, V., Masin, R., and Tarolli, P. (2025). Global impact of seawater intrusion on coastal agriculture. Environ. Res. Lett. 20, 013005. doi: 10.1088/1748-9326/ad9bcd

31. Gorrasi, S., Franzetti, A., Ambrosini, R., Pittino, F., Pasqualetti, M., and Fenice, M. (2021). Spatio-Temporal Variation of the Bacterial Communities along a Salinity Gradient within a Thalassohaline Environment (Saline di Tarquinia Salterns, Italy). Molecules 26, 1338. doi: 10.3390/molecules26051338

32. Gunde-Cimerman, N., Plemenitaš, A., and Oren, A. (2018). Strategies of adaptation of microorganisms of the three domains of life to high salt concentrations. FEMS Microbiology Reviews 42, 353–375. doi: 10.1093/femsre/fuy009

33. Hamilton, T. L., Jones, D. S., Schaperdoth, I., and Macalady, J. L. (2015). Metagenomic insights into S(0) precipitation in a terrestrial subsurface lithoautotrophic ecosystem. Front. Microbiol. 5. doi: 10.3389/fmicb.2014.00756

34. Han, D., and Currell, M. J. (2018). Delineating multiple salinization processes in a coastal plain aquifer, northern China: hydrochemical and isotopic evidence. Hydrol. Earth Syst. Sci. 22, 3473–3491. doi: 10.5194/hess-22-3473-2018

35. Harding, A. (2014). The prehistoric exploitation of salt in Europe. GQ. doi: 10.7306/gq.1164

36. Hino, S., Kawasaki, N., Yamano, N., Nakamura, T., and Nakayama, A. (2023). Effects of particle size on marine biodegradation of poly(l-lactic acid) and poly(ε-caprolactone). Materials Chemistry and Physics 303, 127813. doi: 10.1016/j.matchemphys.2023.127813

37. Houghton, K. M., Fournier, M., and Tschritter, C. (2023). Saltwater Intrusion Impacts Microbial Diversity and Function in Groundwater Ecosystems. TOMICROJ 17, e187428582306190. doi: 10.2174/18742858-v17-230720-2023-2

38. Hu, W., Zheng, N., Zhang, Y., Li, S., Bartlam, M., and Wang, Y. (2024). Metagenomics analysis reveals effects of salinity fluctuation on diversity and ecological functions of high and low nucleic acid content bacteria. Science of The Total Environment 933, 173186. doi: 10.1016/j.scitotenv.2024.173186

39. Hu, W., Zhou, X., Liu, Y., Zhang, Y., and Wang, Y. (2025). Effects of Salinity Fluctuation on Antimicrobial Resistance and Virulence Factor Genes of Low and High Nucleic Acid-Content Bacteria in a Marine Environment. Microorganisms 13, 1710. doi: 10.3390/microorganisms13071710

40. Izabel-Shen, D. (2021). Understanding response of microbial communities to saltwater intrusion through microcosms. Computational and Structural Biotechnology Journal 19, 929–933. doi: 10.1016/j.csbj.2021.01.021

41. Jakab, G., Silye, L., Sümegi, P., Tóth, A., Sümegi, B., Pál, I., et al. (2020). Relict Anthropogenic Ecosystem from the Middle Ages: History of a Salt Marsh from Transylvania (Sic, N Romania). Environmental Archaeology 25, 96–113. doi: 10.1080/14614103.2019.1578547

42. Jasechko, S., Perrone, D., Befus, K. M., Bayani Cardenas, M., Ferguson, G., Gleeson, T., et al. (2017). Global aquifers dominated by fossil groundwaters but wells vulnerable to modern contamination. Nature Geosci 10, 425–429. doi: 10.1038/ngeo2943

43. Jasechko, S., Perrone, D., Seybold, H., Fan, Y., and Kirchner, J. W. (2020). Groundwater level observations in 250,000 coastal US wells reveal scope of potential seawater intrusion. Nat Commun 11, 3229. doi: 10.1038/s41467-020-17038-2

44. Jiao, J., and Post, V. (2019). Coastal Hydrogeology., 1st Edn. Cambridge University Press. doi: 10.1017/9781139344142

45. Jones, D. S., Schaperdoth, I., and Macalady, J. L. (2014). Metagenomic Evidence for Sulfide Oxidation in Extremely Acidic Cave Biofilms. Geomicrobiology Journal 31, 194–204. doi: 10.1080/01490451.2013.834008

46. Kaushal, S. S., Likens, G. E., Mayer, P. M., Shatkay, R. R., Shelton, S. A., Grant, S. B., et al. (2023). The anthropogenic salt cycle. Nat Rev Earth Environ 4, 770–784. doi: 10.1038/s43017-023-00485-y

47. Khalil, M. M., Farag, M. H., Tokunaga, T., Pichler, T., Ismail, E., and Abotalib, A. Z. (2025). Origin and processes of groundwater salinity hotspots in the irrigated Sahara Desert lands of Egypt. Groundwater for Sustainable Development 31, 101520. doi: 10.1016/j.gsd.2025.101520

48. Klindworth, A., Pruesse, E., Schweer, T., Peplies, J., Quast, C., Horn, M., et al. (2013). Evaluation of general 16S ribosomal RNA gene PCR primers for classical and next-generation sequencing-based diversity studies. Nucleic Acids Research 41, e1–e1. doi: 10.1093/nar/gks808

49. Li, C., Gao, X., Li, S., and Bundschuh, J. (2020). A review of the distribution, sources, genesis, and environmental concerns of salinity in groundwater. Environ Sci Pollut Res 27, 41157–41174. doi: 10.1007/s11356-020-10354-6

50. Li, Y., Li, W., Jiang, L., Li, E., Yang, X., and Yang, J. (2024). Salinity affects microbial function genes related to nutrient cycling in arid regions. Front. Microbiol. 15, 1407760. doi: 10.3389/fmicb.2024.1407760

51. Liu, C., Mansoldo, F. R. P., Li, H., Vermelho, A. B., Zeng, R. J., Li, X., et al. (2025). A workflow for statistical analysis and visualization of microbiome omics data using the R microeco package. Nat Protoc. doi: 10.1038/s41596-025-01239-4

52. Liu, W., Jiang, H., Yang, J., and Wu, G. (2018). Gammaproteobacterial Diversity and Carbon Utilization in Response to Salinity in the Lakes on the Qinghai–Tibetan Plateau. Geomicrobiology Journal 35, 392–403. doi: 10.1080/01490451.2017.1378951

53. Louca, S., Parfrey, L. W., and Doebeli, M. (2016). Decoupling function and taxonomy in the global ocean microbiome. Science 353, 1272–1277. doi: 10.1126/science.aaf4507

54. Ma, Z., Gao, L., Sun, M., Liao, Y., Bai, S., Wu, Z., et al. (2022). Microbial Diversity in Groundwater and Its Response to Seawater Intrusion in Beihai City, Southern China. Front. Microbiol. 13, 876665. doi: 10.3389/fmicb.2022.876665

55. Maechler, M., Rousseeuw, P., Struyf, A., and Hubert, M. (1999). cluster: “Finding Groups in Data”: Cluster Analysis Extended Rousseeuw et al. 2.1.8.1. doi: 10.32614/CRAN.package.cluster

56. Mani, K., Salgaonkar, B. B., Das, D., and Bragança, J. M. (2012). Community solar salt production in Goa, India. Aquat. Biosyst. 8, 30. doi: 10.1186/2046-9063-8-30

57. Marien, L., Crabit, A., Dewandel, B., Ladouche, B., Fleury, P., Follain, S., et al. (2023). Salinity spatial patterns in Mediterranean coastal areas: The legacy of historical water infrastructures. Science of The Total Environment 899, 165730. doi: 10.1016/j.scitotenv.2023.165730

58. Martin, M. (2011). Cutadapt removes adapter sequences from high-throughput sequencing reads. EMBnet j. 17, 10. doi: 10.14806/ej.17.1.200

59. Marzano, A. (2024). Marine salt production in the Roman world: The salinae and their ownership. Quaternary Science Reviews 336, 108776. doi: 10.1016/j.quascirev.2024.108776

60. Mastrocicco, M., and Colombani, N. (2021). The Issue of Groundwater Salinization in Coastal Areas of the Mediterranean Region: A Review. Water 13, 90. doi: 10.3390/w13010090

61. Mastrorillo, L., Mazza, R., Manca, F., and Tuccimei, P. (2016). Evidences of different salinization sources in the roman coastal aquifer (Central Italy). J Coast Conserv 20, 423–441. doi: 10.1007/s11852-016-0457-5

62. Melita, M., Amalfitano, S., Preziosi, E., Ghergo, S., Frollini, E., Parrone, D., et al. (2019). Physiological Profiling and Functional Diversity of Groundwater Microbial Communities in a Municipal Solid Waste Landfill Area. Water 11, 2624. doi: 10.3390/w11122624

63. Melita, M., Amalfitano, S., Preziosi, E., Ghergo, S., Frollini, E., Parrone, D., et al. (2023). Redox conditions and a moderate anthropogenic impairment of groundwater quality reflected on the microbial functional traits in a volcanic aquifer. Aquat Sci 85, 3. doi: 10.1007/s00027-022-00899-8

64. Milli, S., D’Ambrogi, C., Bellotti, P., Calderoni, G., Carboni, M. G., Celant, A., et al. (2013). The transition from wave-dominated estuary to wave-dominated delta: The Late Quaternary stratigraphic architecture of Tiber River deltaic succession (Italy). Sedimentary Geology 284-285, 159–180. doi: 10.1016/j.sedgeo.2012.12.003

65. Morelli, C., and Forte, V. (2014). Il Campus Salinarum Romanarum e l’epigrafe dei conductores: Il contesto archeologico. mefra. doi: 10.4000/mefra.2059

66. Mueller, W., Zamrsky, D., Essink, G. O., Fleming, L. E., Deshpande, A., Makris, K. C., et al. (2024). Saltwater intrusion and human health risks for coastal populations under 2050 climate scenarios. Sci Rep 14, 15881. doi: 10.1038/s41598-024-66956-4

67. Narvaez-Montoya, C., Mahlknecht, J., Torres-Martínez, J. A., Mora, A., and Bertrand, G. (2023). Seawater intrusion pattern recognition supported by unsupervised learning: A systematic review and application. Science of The Total Environment 864, 160933. doi: 10.1016/j.scitotenv.2022.160933

68. Nelson, T., Hose, G. C., Dabovic, J., and Korbel, K. L. (2024). Salinity as a major influence on groundwater microbial communities in agricultural landscapes. Marine and Freshwater Research 75, MF23014. doi: 10.1071/MF23014

69. Németh, I., Molnár, S., Vaszita, E., and Molnár, M. (2021). The Biolog EcoPlate^TM^ Technique for Assessing the Effect of Metal Oxide Nanoparticles on Freshwater Microbial Communities. Nanomaterials 11, 1777. doi: 10.3390/nano11071777

70. Oksanen, J., Simpson, G. L., Blanchet, F. G., Kindt, R., Legendre, P., Minchin, P. R., et al. (2001). vegan: Community Ecology Package. 2.7–2. doi: 10.32614/CRAN.package.vegan

71. Ongetta, S., Mohan Viswanathan, P., Sabarathinam, C., Ramasamy, N., and Kuek, C. (2022). Delineation of highland saline groundwater sources in Ba’kelalan region of NE Borneo to improve the salt-making production using geochemical and geophysical approaches. Chemosphere 307, 135721. doi: 10.1016/j.chemosphere.2022.135721

72. Passarella, G., Masciale, R., Menichini, M., Doveri, M., and Portoghese, I. (2025). Decoding Salinization Dynamics in Mediterranean Coastal Aquifers: A Case Study from a Wetland in Southern Italy. Environments 12, 227. doi: 10.3390/environments12070227

73. Payler, S. J., Biddle, J. F., Sherwood Lollar, B., Fox-Powell, M. G., Edwards, T., Ngwenya, B. T., et al. (2019). An Ionic Limit to Life in the Deep Subsurface. Front. Microbiol. 10, 426. doi: 10.3389/fmicb.2019.00426

74. Percak-Dennett, E. M., and Roden, E. E. (2014). Geochemical and Microbiological Responses to Oxidant Introduction into Reduced Subsurface Sediment from the Hanford 300 Area, Washington. Environ. Sci. Technol. 48, 9197–9204. doi: 10.1021/es5009856

75. Quast, C., Pruesse, E., Yilmaz, P., Gerken, J., Schweer, T., Yarza, P., et al. (2012). The SILVA ribosomal RNA gene database project: improved data processing and web-based tools. Nucleic Acids Research 41, D590–D596. doi: 10.1093/nar/gks1219

76. Rath, K. M., and Rousk, J. (2015). Salt effects on the soil microbial decomposer community and their role in organic carbon cycling: A review. Soil Biology and Biochemistry 81, 108–123. doi: 10.1016/j.soilbio.2014.11.001

77. Renau-Pruñonosa, A., Esteller, M. V., Aroba, J., Grande, J. A., Morell, I., De La Torre, M. L., et al. (2025). Identification of salinization processes in coastal aquifers using a fuzzy logic and data mining based methodology: study case in a Mediterranian coastal aquifer (Spain). Environ Earth Sci 84, 109. doi: 10.1007/s12665-024-12006-1

78. Risley, C. A., Tamalavage, A. E., Van Hengstum, P. J., and Labonté, J. M. (2022). Subsurface Microbial Community Composition in Anchialine Environments Is Influenced by Original Organic Carbon Source at Time of Deposition. Front. Mar. Sci. 9, 872789. doi: 10.3389/fmars.2022.872789

79. Robinson, H. K., Hasenmueller, E. A., and Chambers, L. G. (2017). Soil as a reservoir for road salt retention leading to its gradual release to groundwater. Applied Geochemistry 83, 72–85. doi: 10.1016/j.apgeochem.2017.01.018

80. Rossmassler, K., Hanson, T. E., and Campbell, B. J. (2016). Diverse sulfur metabolisms from two subterranean sulfidic spring systems. FEMS Microbiology Letters 363, fnw162. doi: 10.1093/femsle/fnw162

81. Sang, S., Jiang, H., Gan, J., and Ke, C. (2025). Microbial community dynamics across salinity gradients in coastal aquifers: Linking hydrogeochemical variability to prokaryotic diversity in a seawater-intruded aquifer of the Pearl River Delta, China. Marine Environmental Research 211, 107471. doi: 10.1016/j.marenvres.2025.107471

82. Segata, N., Izard, J., Waldron, L., Gevers, D., Miropolsky, L., Garrett, W. S., et al. (2011). Metagenomic biomarker discovery and explanation. Genome Biol 12, R60. doi: 10.1186/gb-2011-12-6-r60

83. Seibert, S. L., Greskowiak, J., Oude Essink, G. H. P., and Massmann, G. (2024). Understanding Climate Change and Anthropogenic Impacts on the Salinization of Low-Lying Coastal Groundwater Systems. Earth’s Future 12, e2024EF004737. doi: 10.1029/2024EF004737

84. Selak, L., Meier, D. V., Marinović, M., Čačković, A., Kajan, K., Pjevac, P., et al. (2025). Salinization alters microbial methane cycling in freshwater sediments. Environmental Microbiome 20, 73. doi: 10.1186/s40793-025-00739-w

85. Selvakumar, S., Chandrasekar, N., Srinivas, Y., Selvam, S., Kaliraj, S., Magesh, N. S., et al. (2022). Hydrogeochemical processes controlling the groundwater salinity in the coastal aquifers of Southern Tamil Nadu, India. Marine Pollution Bulletin 174, 113264. doi: 10.1016/j.marpolbul.2021.113264

86. USEPA (2000). Guidance for data quality assessment. Practical methods for data analysis. Office of Environmental Information. EPA QA/G-9, QA00 Version Washington, DC.

87. Valle, A. (2023). Studio idrogeochimico del sito aeroportuale di Fiumicino finalizzato alla definizione del modello idrogeologico concettuale. Roma, Italia: Università degli Studi di Roma, Dipartimento di Scienze della Terra.

88. Vicari, F., Randazzo, S., López, J., Fernández De Labastida, M., Vallès, V., Micale, G., et al. (2022). Mining minerals and critical raw materials from bittern: Understanding metal ions fate in saltwork ponds. Science of The Total Environment 847, 157544. doi: 10.1016/j.scitotenv.2022.157544

89. Wang, H., Herrmann, M., Schroeter, S. A., Zerfaß, C., Lehmann, R., Lehmann, K., et al. (2025). Groundwater microbiomes balance resilience and vulnerability to hydroclimatic extremes. Commun Earth Environ 6, 683. doi: 10.1038/s43247-025-02680-9

90. Wang, J., Liu, Y., Ma, Y., Wang, X., Zhang, B., Zhang, G., et al. (2023). Research progress regarding the role of halophilic and halotolerant microorganisms in the eco-environmental sustainability and conservation. Journal of Cleaner Production 418, 138054. doi: 10.1016/j.jclepro.2023.138054

91. Wang, J., Zhang, Y., Ding, Y., Zhang, Y., Xu, W., Zhang, X., et al. (2024). Adaptive characteristics of indigenous microflora in an organically contaminated high salinity groundwater. Chemosphere 349, 140951. doi: 10.1016/j.chemosphere.2023.140951

92. Watson, R., and McKillop, H. (2019). A Filtered Past: Interpreting Salt Production and Trade Models from Two Remnant Brine-Enrichment Mounds at the Ancient Maya Paynes Creek Salt Works, Belize. Journal of Field Archaeology 44, 40–51. doi: 10.1080/00934690.2018.1557993

93. Weinstein, M. M., Prem, A., Jin, M., Tang, S., and Bhasin, J. M. (2019). FIGARO: An efficient and objective tool for optimizing microbiome rRNA gene trimming parameters. doi: 10.1101/610394

94. Wen, S., Liang, S., Pang, G., Shan, Q., Ye, Y., Zhang, J., et al. (2025). Hydrochemical Characteristics and Evolution Mechanisms of Shallow Groundwater in the Alluvial–Coastal Transition Zone of the Tangshan Plain, China. Water 17, 2810. doi: 10.3390/w17192810

95. Werner, A. D., Bakker, M., Post, V. E. A., Vandenbohede, A., Lu, C., Ataie-Ashtiani, B., et al. (2013). Seawater intrusion processes, investigation and management: Recent advances and future challenges. Advances in Water Resources 51, 3–26. doi: 10.1016/j.advwatres.2012.03.004

96. West, A. G., Boey, J. S., Tee, H. S., and Handley, K. M. (2025). Salinity-driven niche differentiation within the aquatic Luna-1 subcluster. ISME Communications 5, ycaf122. doi: 10.1093/ismeco/ycaf122

97. Wu, Y., and Wang, Y. (2014). Geochemical evolution of groundwater salinity at basin scale: a case study from Datong basin, northern China. Environ. Sci.: Processes Impacts 16, 1469. doi: 10.1039/c4em00019f

98. Xamxidin, M., Zhang, X., Zheng, G., Chen, C., and Wu, M. (2025). Metagenomics-assembled genomes reveal microbial metabolic adaptation to athalassohaline environment, the case Lake Barkol, China. Front. Microbiol. 16, 1550346. doi: 10.3389/fmicb.2025.1550346

99. Xiu, W., Lloyd, J., Guo, H., Dai, W., Nixon, S., Bassil, N. M., et al. (2020). Linking microbial community composition to hydrogeochemistry in the western Hetao Basin: Potential importance of ammonium as an electron donor during arsenic mobilization. Environment International 136, 105489. doi: 10.1016/j.envint.2020.105489

100. Xu, X., Xiong, G., Chen, G., Fu, T., Yu, H., Wu, J., et al. (2021). Characteristics of coastal aquifer contamination by seawater intrusion and anthropogenic activities in the coastal areas of the Bohai Sea, eastern China. Journal of Asian Earth Sciences 217, 104830. doi: 10.1016/j.jseaes.2021.104830

101. Yang, J., Li, W., Teng, D., Yang, X., Zhang, Y., and Li, Y. (2022). Metagenomic Insights into Microbial Community Structure, Function, and Salt Adaptation in Saline Soils of Arid Land, China. Microorganisms 10, 2183. doi: 10.3390/microorganisms10112183

102. Zhang, X., Qi, L., Li, W., Hu, B. X., and Dai, Z. (2021). Bacterial community variations with salinity in the saltwater-intruded estuarine aquifer. Science of The Total Environment 755, 142423. doi: 10.1016/j.scitotenv.2020.142423

103. Zheng, T., Deng, Y., Wang, Y., Jiang, H., O’Loughlin, E. J., Flynn, T. M., et al. (2019). Seasonal microbial variation accounts for arsenic dynamics in shallow alluvial aquifer systems. Journal of Hazardous Materials 367, 109–119. doi: 10.1016/j.jhazmat.2018.12.087

104. Zhi, C., Hu, X., Yang, F., Huang, X., Chen, H., Chen, L., et al. (2024). Unraveling microbial community variation along a salinity gradient and indicative significance to groundwater salinization in the coastal aquifer. Journal of Hydrology 642, 131893. doi: 10.1016/j.jhydrol.2024.131893

105. Zoppini, A., Bongiorni, L., Ademollo, N., Patrolecco, L., Cibic, T., Franzo, A., et al. (2020). Bacterial diversity and microbial functional responses to organic matter composition and persistent organic pollutants in deltaic lagoon sediments. Estuarine, Coastal and Shelf Science 233, 106508. doi: 10.1016/j.ecss.2019.106508

